# Wnt activator FOXB2 drives prostate cancer neuroendocrine differentiation

**DOI:** 10.1101/564625

**Authors:** Lavanya Moparthi, Giulia Pizzolato, Stefan Koch

## Abstract

The Wnt signaling pathway is of paramount importance for development and disease. However, the tissue-specific regulation of Wnt pathway activity remains incompletely understood. Here we identify FOXB2, an uncharacterized forkhead box family transcription factor, as a potent activator of Wnt signaling in normal and cancer cells. Mechanistically, FOXB2 induces the non-classical Wnt ligand WNT7B, which increases TCF/LEF-dependent transcription without activating LRP6 or β-catenin. Proximity ligation and RNA interference identified YAP1, JUN, and DDX5 as transcriptional co-regulators required for FOXB2-dependent Wnt activation. Although FOXB2 expression is limited in adults, it is induced in select cancers, particularly advanced prostate cancer. RNA-seq data analysis suggests that FOXB2/WNT7B expression in prostate cancer is associated with a transcriptional program that favors neuronal differentiation and decreases recurrence-free survival. Consistently, FOXB2 is induced during neuroendocrine transformation of LNCaP prostate carcinoma cells, and conversely, FOXB2 overexpression is sufficient to induce their differentiation. Our results suggest that FOXB2 is a tissue-specific Wnt enhancer that promotes prostate cancer malignant transformation.

## INTRODUCTION

The Wnt pathway is a major homeostatic signaling cascade in development and stem cell homeostasis (Koch, 2017; MacDonald et al., 2009; Nusse and Clevers, 2017). In the canonical or β-catenin-dependent signaling branch, secreted Wnt ligands engage a transmembrane receptor complex consisting of Frizzled family core and LRP5/6 co-receptors, to inhibit a multiprotein β-catenin destruction complex. Consequently, cytosolic β-catenin is relieved from constitutive proteasomal degradation, and induces the transcription of target genes through association with TCF/LEF family transcription factors.

Specificity of the Wnt signaling output is achieved primarily via the differential expression of a wide range of Wnt ligands and receptors, which exert overlapping but non-redundant functions (MacDonald et al., 2009). Interestingly, several Wnt molecules act downstream of β-catenin, by synergizing with canonical Wnt ligands. For example, WNT7B elicits limited pathway activation on its own, as evidenced by its inability to induce LRP6 phosphorylation and β-catenin stabilization to any substantial degree (Alok et al., 2017). However, WNT7B strongly co-operates with other ligands, primarily Wnt1, in driving TCF/LEF-dependent gene transcription. The mechanism of WNT7B-dependent pathway activation is unclear, but it requires additional co-receptors, namely RECK and GPR124 (Alok et al., 2017; Vallon et al., 2018; Vanhollebeke et al., 2015). The expression of WNT7B and its co-receptors is largely restricted to specific tissues, especially the developing brain, where they contribute to blood-brain barrier formation and maintenance through activation of Wnt/β-catenin signaling (Kuhnert et al., 2010; Zhou and Nathans, 2014). Additionally, increased expression of *WNT7B* and subsequent Wnt pathway activation have been observed in cancers, including gliomas and pancreatic cancer (Arensman et al., 2013; Griveau et al., 2018), although it is unclear what causes *WNT7B* induction in these diseases.

In this study, we identify the uncharacterized forkhead box (FOX) transcription factor FOXB2 as a potent activator of WNT7B-dependent TCF signaling. Although *FOXB2* expression is largely restricted to the developing brain (Kaestner et al., 1996), we find that it is induced in advanced prostate cancer. Most prostate tumors initially progress slowly, but clonal evolution of cancer cells eventually results in androgen resistance and neuroendocrine differentiation, which is associated with treatment failure and exceptionally poor prognosis (Parimi et al., 2014). Chronic Wnt pathway activation is a key driver of malignant prostate cancer progression (Murillo-Garzón and Kypta, 2017). However, in contrast to e.g. colorectal cancer, activating pathway mutations are relatively infrequent in prostate cancer, and sustained Wnt signaling is thought to be maintained via tissue-specific pathway activators, including WNT7B (Murillo-Garzón and Kypta, 2017; Zheng et al., 2013). Thus, our identification of FOXB2 as major WNT7B regulator may have important implications for developmental and cancer biology.

## MATERIALS AND METHODS

### Cell culture

Authenticated 293T, HCT116, SW48, LNCaP, L, and L/Wnt3a cells were obtained from the German Collection of Microorganisms and Cell Cultures (DSMZ, Braunschweig, Germany), the American Type Culture Collection (ATCC, Manassas, USA) and the European Collection of Authenticated Cell Cultures (ECACC, Salisbury, UK). 293T ΔLRP6 were generated by CRISPR/Cas9 mediated gene editing, essentially as described (Berger et al., 2017). Briefly, an enhanced specificity Cas9 vector with gRNA ATTATTGTCCCCCGATGGGC was transiently transfected into 293T cells, and cell clones were obtained by limiting dilution. 293T ΔCTNNB1 and ΔTCF/LEF have been described elsewhere (Doumpas et al., 2019). Cells were maintained at 5% CO_2_ / 37°C in RPMI 1640 (LNCaP) or DMEM (all other cells) with 10% fetal bovine serum (FBS), 2 mM glutamine, and 1% penicillin/streptomycin. All experiments were performed using low passage cells from confirmed mycoplasma-free frozen stocks.

Wnt3a and control conditioned media were collected from stably transfected L cells, following the supplier’s guidelines. R-spondin 3 conditioned media were generated by transient transfection of Rspo3ΔC (Ohkawara et al., 2011) into 293T cells.

### Molecular cloning

Expression constructs were generated by restriction cloning of full-length cDNA from an in-house intestinal epithelial cell library into a pCS2+ vector with N-terminal Flag or V5 tag. For stable expression, cDNAs were subcloned into a pcDNA3 vector, and transfectants were isolated by G418 selection. Point and truncation mutants were generated by restriction cloning and PCR-based mutagenesis, using high fidelity Q5 polymerase (New England Biolabs, Ipswich, US). All plasmids were validated by partial sequencing.

### Plasmid and siRNA transfection

Cells were transfected by calcium phosphate precipitation (293T only), or lipofection using Lipofectamine 3000 (LNCaP) or 2000 (all other cells) (Thermo Fisher, Waltham, USA). siRNAs were purchased from Integrated DNA Technologies (IDT, Skokie, USA) and Thermo Fisher. Detailed plasmid information can be found in Supplemental Table 1.

### Antibodies and reagents

Primary antibodies were purchased from the following companies. Sigma Aldrich (St. Louis, USA): mouse anti-Flag M2 (F3165), rabbit anti-FOXB2 (HPA067947); Novus Biologicals (Centennial, USA): mouse anti-NSE (NBP2-47635); Antibodies-online (Aachen, Germany): rabbit anti-β-catenin (ABIN1881238); Cell Signaling Technology (Danvers, USA): rabbit anti-LRP6 (3395), rabbit anti-Active β-catenin (8814), rat anti-HSP70 (4873); Genetex (Irvine, USA): rabbit anti-LRP6 Sp1490 (GTX62033); Santa Cruz Biotechnology (Dallas, USA): mouse anti-Transferrin receptor (sc-65882), mouse anti-LRP6 (sc-25317), mouse anti-β-catenin (sc-133240). Chemicals and inhibitors were from Sigma Aldrich and Cayman Chemicals (Ann Arbor, USA).

### Luciferase assays

The TOPflash β-catenin/TCF reporter assay was performed essentially as described (Veeman et al., 2003). The WNT7B reporter was generated by replacing the TOPflash TCF binding domains with a 1kb enhancer region from the human WNT7B gene. Luciferase activity was measured using a Firefly luciferase glow assay kit (Thermo Fisher). Data were normalized to the average intensity of identically treated FOPflash control, performed in triplicate.

### Quantitative real-time PCR

qPCR was performed using standard protocols. In brief, RNA was extracted using a Qiagen RNeasy Mini kit (Hilden, Germany), and reverse transcribed with a Thermo Fisher cDNA synthesis kit. cDNA was amplified using validated custom primers, with SYBR green dye. Data were acquired on a Bio-Rad CFX96 Touch thermocycler (Hercules, USA), and normalized to *HPRT1* control.

### Immunoblotting

Immunoblotting was performed using standard protocols. In brief, proteins were extracted in NP-40 containing lysis buffer with protease inhibitors, and boiled in Laemmli sample buffer. Subcellular fractionation was done with the REAP method (Suzuki et al., 2010). Samples were separated on polyacrylamide gels, transferred onto nitrocellulose membranes, and incubated in blocking buffer. Primary antibodies were detected using near infrared (NIR) fluorophore-labeled secondary antibodies. Blots were scanned on a LI-COR CLx imager (Lincoln, USA). Consumables were purchased from Bio-Rad and LI-COR.

### Co-immunoprecipitation

Co-IP was performed essentially as before (Koch et al., 2015), with minor modifications. Briefly, constructs were expressed in HCT116 cells for 48 hours, and nuclei were isolated using the REAP method. Nuclei were resuspended in IP lysis buffer (TBS pH 7.5, 1mM MgCl_2_, 2mM β-mercaptoethanol, 10mM sodium pyrophosphate, 1% v/v Triton X-100, protease inhibitors) and sonicated. Samples were precleared with protein A/G beads (Santa Cruz Biotechnology) for 1 hour, and incubated with anti-Flag antibody for 2 hours at 4°C with end-over-end rotation. Protein A/G beads were added, and samples were incubated overnight at 4°C with end-over-end rotation. After repeated washes in IP wash buffer (TBS pH 7.5, 2mM β-mercaptoethanol, 1% v/v Triton X-100), beads were boiled in Laemmli sample buffer and analyzed by immunoblotting. 5% of the initial nuclear lysate was saved for input control.

### Immunocytochemistry

Immunocytochemistry was performed essentially as before (Koch et al., 2015). In brief, cells were fixed with 4% paraformaldehyde, permeabilized with Triton X-100 containing buffer, and blocked with bovine serum albumin. Primary antibodies were detected using fluorophore-labeled secondary antibodies (Thermo Fisher). Samples were mounted in Prolong Glass Antifade (Thermo Fisher) with Hoechst 33342 or NucBlue nuclear counterstain. Images were acquired on a Zeiss LSM 800 confocal microscope with Airyscan (Oberkochen, Germany), or a Nikon E800 epifluorescence microscope (Amstelveen, Netherlands). Fluorescence intensity on a per-cell level was measured in ImageJ 1.52i (National Institutes of Health, USA).

### BioID and mass spectrometry

The BioID assay was performed essentially as described (Roux et al., 2012). N-terminal BirA-FOXB2 and BirA plasmids were transfected into 293T cells using Lipofectamine 2000. After 6 h of transfection, cells were treated with 50 μM biotin, and incubated for 24 h at 37°C, 5% CO2. Cells were lysed in RIPA buffer for 1 h at 4 °C with end-over-end rotation. Pre-washed streptavidin beads (GE Healthcare, USA) were added to the cell lysate and incubated for 3 h at 4°C with end-over-end rotation. The beads were washed four times with 50 mM ammonium bicarbonate (NH_4_HCO_3_). Enzymatic digestion was carried out by adding 100 μl of freshly prepared 10 ng/μl proteomics grade trypsin (Thermo Fisher) in 50 mM NH_4_HCO_3_ to the beads, and samples were incubated overnight at 37°C with end-over-end rotation. The digested samples were dried by vacuum centrifugation.

BioID experiments were performed in triplicates using an Easy nano LC II HPLC interfaced with a nanoEasy spray ion source (Thermo Fisher) connected to an Orbitrap Velos Pro mass spectrometer (Thermo Fisher). The peptides were loaded on a NS-MP-10 Biosphere C18 column 2 cm (100 μm inner diameter, packed with 5 μm resin) and the chromatographic separation was performed at RT on a 10.1 cm (75 μm inner diameter) NS-AC-10-C18 column packed with 5 μm resin (NanoSeperations, Netherlands). The nano HPLC was operating at 300 nl/min flow rate with a linear gradient of solvent B (0.1% (v/v) formic acid in acetonitrile) in solvent A (0.1% (v/v) formic acid in water) for 60 min.

An MS scan (380–2000 m/z) was recorded in the FTMS mass analyzer set at a resolution of 30,000. The MS was followed by top speed data-dependent collision-induced dissociation MS/MS scans on multiply charged ions. The general mass spectrometric conditions were as follows: spray voltage, 3 kV; no sheath or auxiliary gas flow; S-lens 65.7%; ion transfer tube temperature, 225°C. Collision-induced dissociation was applied with 35% of energy, and MS/MS spectra were recorded in the ion trap at a resolution of 15,000.

Raw data were processed by Proteome Discoverer 2.0 (Thermo Fisher Scientific) searching the UniProt database with Sequest HT search engine. The search parameters were: Taxonomy: Homo sapiens; Enzymes: trypsin with one missed cleavage, no variable Modifications; Peptide Mass Tolerance: 10 ppm; MS/MS Fragment Tolerance: 0.5 Da. Quantification of the analyzed data was performed with Scaffold 4 (Proteome Software, Portland, USA) using total spectral count.

### In-situ proximity ligation

Proximity ligation was performed using Duolink PLA assay reagents according to the supplier’s guidelines (Sigma Aldrich). In brief, cells were transiently transfected with V5-FOXB2 and Flag-tagged proteins of interest, and antigens were detected with rabbit anti-FOXB2 and mouse anti-Flag antibodies. Following addition of matched PLA probes, rolling circle amplification proceeded for 90 minutes at 37°C. Identically transfected cells were processed for regular immunocytochemistry using the same antibodies, to control for protein expression and localization. An empty eGFP vector at 1/10^th^ the total plasmid amount was added in PLA assays to identify transfected cells.

### Public dataset analyses

The results in this study are in large part based upon data generated by the TCGA Research Network (http://cancergenome.nih.gov/). Dataset analyses were performed primarily in the cBioPortal for Cancer Genomics (Gao et al., 2013) and GEPIA (Tang et al., 2017), with additional analyses and visualization done in R 3.5.2 (R Foundation for Statistical Computing). For genome-wide correlation analysis, the Spearman correlation values of all genes versus FOXB2 or WNT7B were extracted from dataset Prostate Adenocarcinoma (TCGA, Provisional) in the cBioPortal, and co-correlated in R using the *cor.test* function. For *WNT7B* expression in prostate cancer, transcript levels were determined in all tumors with mRNA upregulation versus unaltered samples. Groups were compared with an unpaired, two-tailed Student’s t-test. All survival analyses were performed in GEPIA 2, based on median gene expression. Gene ontology analyses were performed in DAVID (Huang et al., 2007), using the Biological Process Direct function. *p* values were corrected using the Benjamini-Hochberg FDR method.

### Reproducibility and experimental controls

All experimental data are representative of two or more independent experiments with comparable results. Individual data points depict biological replicates. All experiments were performed with appropriate controls, e.g. empty vector, drug vehicle, non-targeting siRNA, and control conditioned media, at identical concentrations. Data and reagents generated in this study will be shared by the lead contact upon reasonable request.

## RESULTS

### FOXB2 activates Wnt/TCF signaling

The context-dependent regulation of canonical Wnt signaling is incompletely understood. In an effort to discover novel pathway regulators, we performed a gain-of-function screen by co-expressing Flag-tagged proteins of interest with a β-catenin/TCF luciferase reporter (TOPflash) in HEK 293T cells. We identified the uncharacterized transcription factor FOXB2 as a candidate Wnt activator (Fig. 1A, B). FOXB2 strongly promoted TOPflash activity, at least to the same extent as the known Wnt activator FOXQ1 (Peng et al., 2015), and potently synergized with the pathway agonists Wnt3a/R-spondin 3 (Fig. 1B). Moreover, FOXB2 induced TOPflash activity in HCT116 and SW48 colorectal cancer cells, which harbor activating β-catenin mutations and are thus largely unresponsive to further pathway activation (Fig. 1C). Of note, the most closely related FOX family member, FOXB1, as well as other tested FOX proteins, did not activate Wnt signaling in these assays (Fig. S1A, and data not shown).

**Figure 1:**
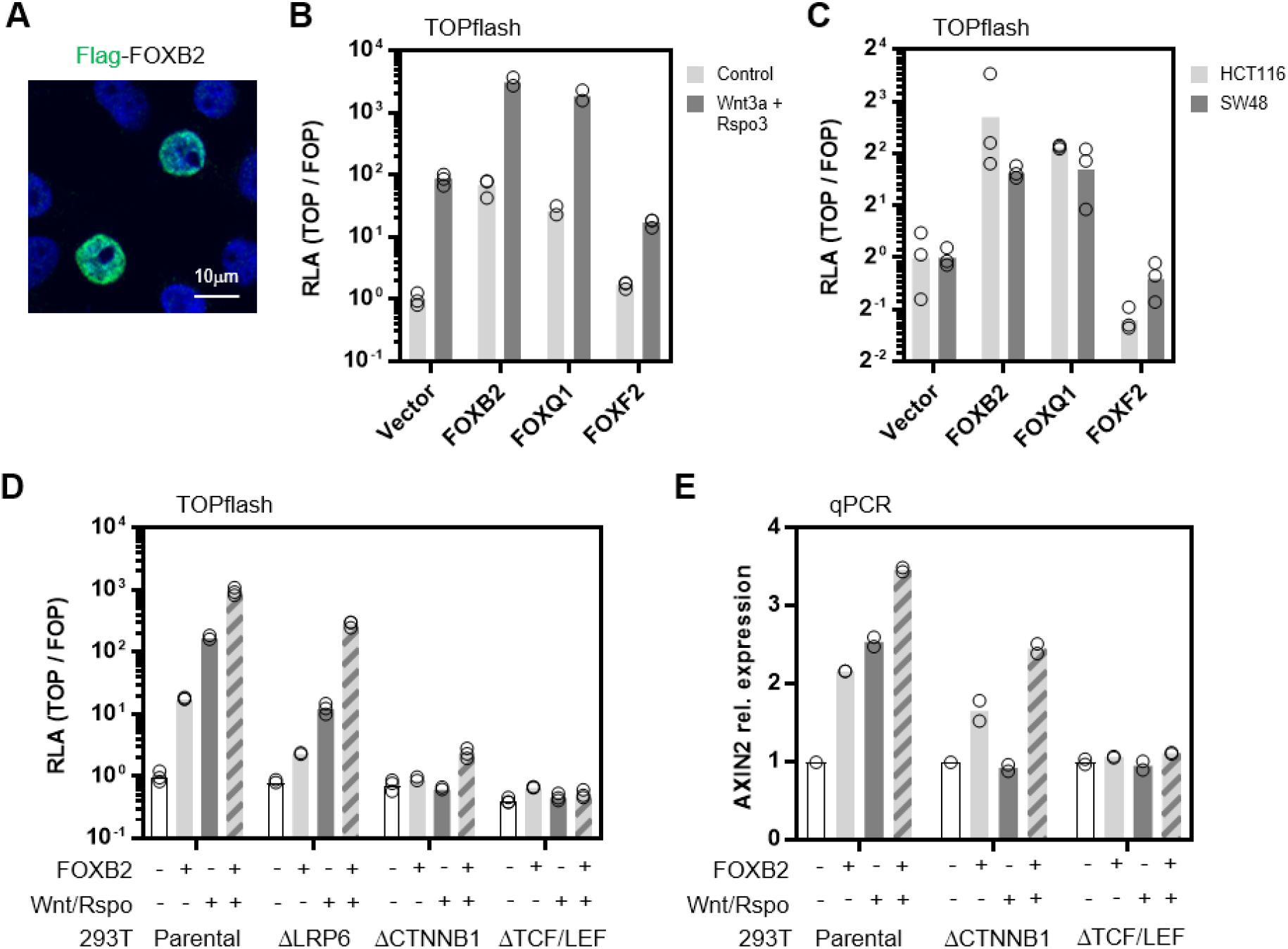
FOXB2 is a Wnt/TCF activator. (**A**) Over-expression in 293T cells showed nuclear FOXB2 localization. (**B**) β-catenin/TCF luciferase reporter assay (TOPflash) in 293T cells. Expression of the indicated Flag-tagged proteins showed that FOXB2 activates Wnt signaling, and synergizes with Wnt3a/R-spondin (Rspo) 3 conditioned media in pathway activation. FOXQ1 and FOXF2 were included as positive and negative controls, respectively. RLA, relative luciferase activity. (**C**) FOXB2 activated TOPflash in HCT116 and SW48 colorectal cancer cells with activating β-catenin mutation. (**D**) Epistasis assay in normal and gene edited 293T. Loss of LRP6, β-catenin (CTNNB1), or all TCF/LEF transcription factors strongly attenuated FOXB2-dependent reporter activation. (**E**) Quantitative real-time PCR (qPCR) in 293T cells. FOXB2 induced *AXIN2* expression in synergy with Wnt3a/Rspo 3. This effect was also observed in β-catenin-deficient, but not TCF/LEF-deficient cells.

FOX transcription factors control Wnt signaling through various mechanisms, such as the regulation of β-catenin nuclear shuttling and stability (Higashimori et al., 2018; Peng et al., 2015; Zhang et al., 2011). To elucidate the signaling mode of FOXB2, we first performed an epistasis assay in gene-edited 293T cells with genetic deletion of Wnt co-receptor LRP6, β-catenin, or all TCF/LEF transcription factors (Doumpas et al., 2019) (Fig. S1B, C, and Fig. 1D). Loss of LRP6 was sufficient to reduce FOXB2-induced TOPflash activation by approximately 90%, while deletion of β-catenin and TCF/LEF essentially blocked reporter activity. Moreover, in contrast to e.g. FOXQ1 and FOXM1 (Bagati et al., 2017; Zhang et al., 2011), FOXB2 did not bind β-catenin or co-localize with nuclear β-catenin (Fig. S1D, E). Since TOPflash is an artificial Wnt reporter, we also tested the regulation of the prototypical Wnt target gene *AXIN2* by qPCR (Fig. 1E). Consistent with results from the TOPflash assays, FOXB2 induced *AXIN2* expression alone as well as in synergy with Wnt3a/R-spondin 3. Surprisingly, partial *AXIN2* induction by FOXB2 was also observed in β-catenin deficient cells, and Wnt3a/R-spondin 3 synergy was retained despite no effect of these ligands by themselves. Collectively, these data identify FOXB2 as a novel Wnt pathway activator upstream of β-catenin.

### FOXB2 induces WNT7B to activate TCF-dependent transcription

The most likely explanation for the above observations is that FOXB2 activates Wnt signaling by inducing one or more canonical Wnt ligands. To test this hypothesis, we first treated FOXB2-transfected 293T cells with porcupine inhibitor LGK974, which blocks the release of endogenous Wnts (Fig. 2A). LGK974 strongly attenuated FOXB2-induced Wnt signaling. This effect was more pronounced in cells with additional exogenous R-spondin 3 compared to cells treated with Wnt3a conditioned media, suggesting that FOXB2 induces Wnt ligands rather than R-spondins. Consistently, iCRT14, which inhibits β-catenin/TCF interaction, also reduced FOXB2-dependent TOPflash activation, and this effect was less pronounced in the presence of exogenous Wnt3a (Fig. S2).

**Figure 2:**
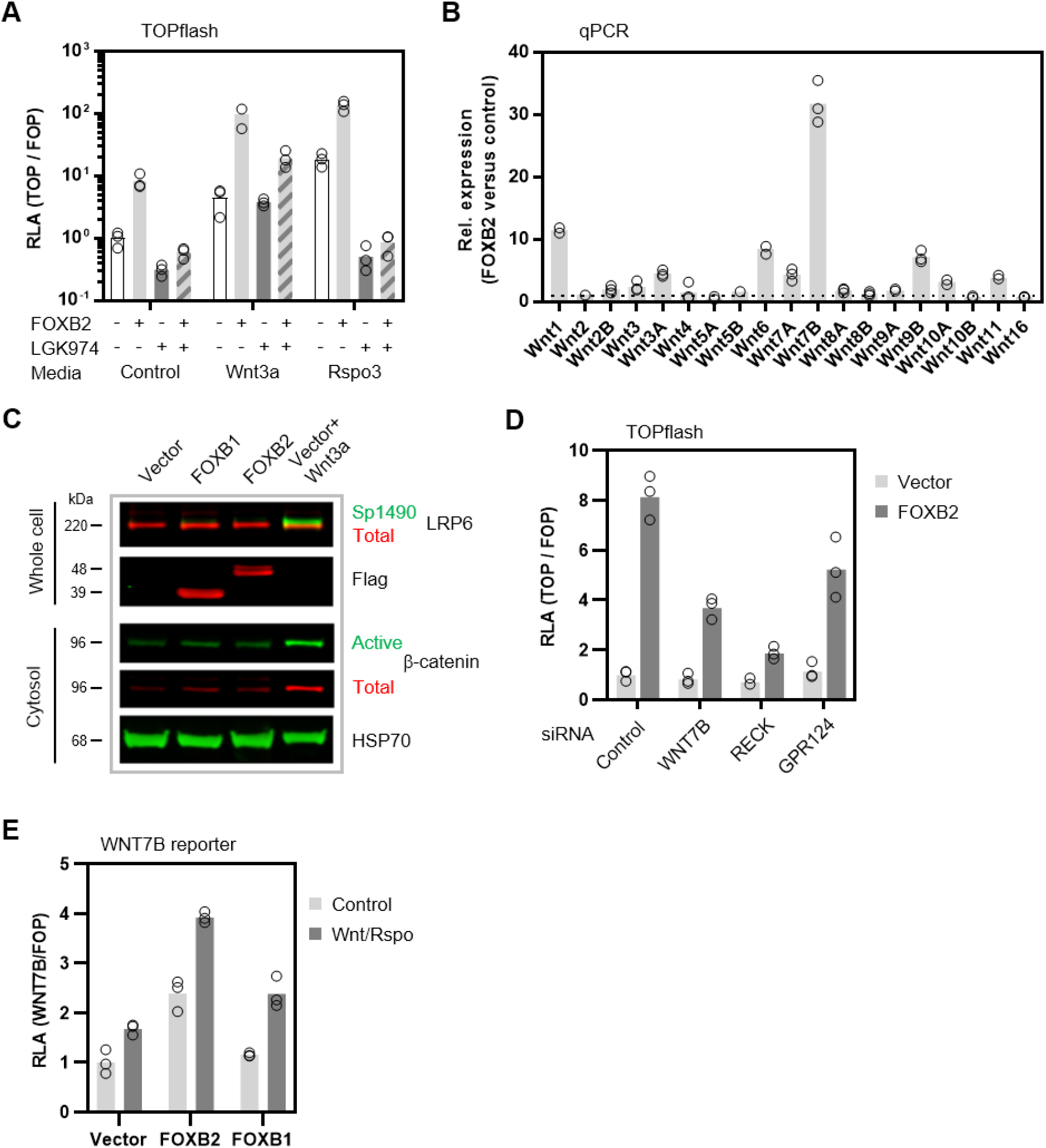
FOXB2 activates TCF signaling via WNT7B. (**A**) Treatment with Wnt secretion inhibitor LGK974 (10μM) strongly attenuated FOXB2-dependent TOPflash activation in 293T cells, particularly in the presence of exogenous R-spondin 3. (**B**) qPCR of all 19 Wnt ligands in 293T cells. FOXB2 strongly induced multiple Wnts. Data are displayed as fold change compared to empty vector control. (**C**) Immunoblot of 293T whole cell lysates and corresponding cytosolic extracts. FOXB2 did not activate LRP6 or stabilize β-catenin. Where indicated, cells were treated with Wnt3a conditioned media for 6 hours. (**D**) Depletion of WNT7B or its co-receptors RECK and GPR124 by RNA interference attenuated FOXB2-dependent TOPflash activation in 293T cells. Note that these siRNAs had essentially no effect on basal Wnt signaling. (**E**) Luciferase-based WNT7B enhancer assay in 293T cells. FOXB2, but not FOXB1, activated the WNT7B reporter in synergy with Wnt3a/R-spondin 3.

To explore these observations further, we examined the regulation of all 19 Wnt ligands by FOXB2 (Fig. 2B). Expression of FOXB2 in 293T cells strongly induced multiple Wnts, particularly WNT1, WNT6, WNT7B, and WNT9B. Since several of these ligands can activate the canonical Wnt signaling branch, we next examined LRP6 phosphorylation and β-catenin stabilization, which are early hallmarks of pathway activation (Fig. 2C). Somewhat surprisingly, FOXB2 did not activate Wnt signaling through LRP6 and β-catenin, despite strong activity at the transcriptional level. We note, however, that this exact effect has recently been reported for some non-classical Wnt ligands, particularly WNT7B (Alok et al., 2017), which was most strongly induced by FOXB2. Thus, to determine if WNT7B is the major mediator of FOXB2-induced Wnt signaling, we interrupted this pathway branch by RNA interference (Fig. 2D). Indeed, depletion of WNT7B or its co-receptors RECK and GPR124 strongly attenuated TOPflash activation by FOXB2. Finally, to test whether FOXB2 directly regulates WNT7B, we generated a luciferase reporter construct containing an intronic *WNT7B* enhancer region (Fig. 2E). FOXB2, but not FOXB1, activated this reporter, and again synergized with Wnt3a/R-spondin 3. We conclude that FOXB2 activates Wnt/TCF independently of β-catenin, by directly engaging a non-classical WNT7B/RECK/GPR124 signaling module.

### Protein-DNA and protein-protein interaction shape FOXB2-dependent Wnt signaling

The DNA-binding forkhead box is highly conserved across FOX family proteins (Li and Tucker, 1993) (Figure S3). Nonetheless, despite exceedingly high sequence similarity between e.g. the FOXB1 and FOXB2 forkhead domain (94% amino acid identity), only FOXB2 promotes Wnt signaling. Thus, the control of specific target genes likely requires interaction of FOXB2 with additional transcription regulators, which is consistent with the regulation of other FOX proteins (Myatt and Lam, 2007). To explore the structural requirements for FOXB2-induced Wnt signaling, we first generated a series of truncations and point mutants (Fig. S4A). Importantly, all truncation constructs exhibited strongly reduced activity in TOPflash assay, including deletion mutants missing the unique central and C-terminal regions (Fig. 3A). Moreover, a FOXB2 point mutant that is unable to bind DNA (P14A/P15A, see (Li and Tucker, 1993)) was completely inert in this assay. In contrast, mutation of a putative engrailed homology (EH1) motif (G277A) had no effect on TOPflash activation. Based on these results, we tested the ability of select FOXB2 mutants to induce *WNT7B* expression (Fig. 3B). Consistent with the TOPflash data, *WNT7B* induction was essentially blocked in FOXB2 constructs lacking either the conserved N-terminal or unique C-terminal domain.

**Figure 3:**
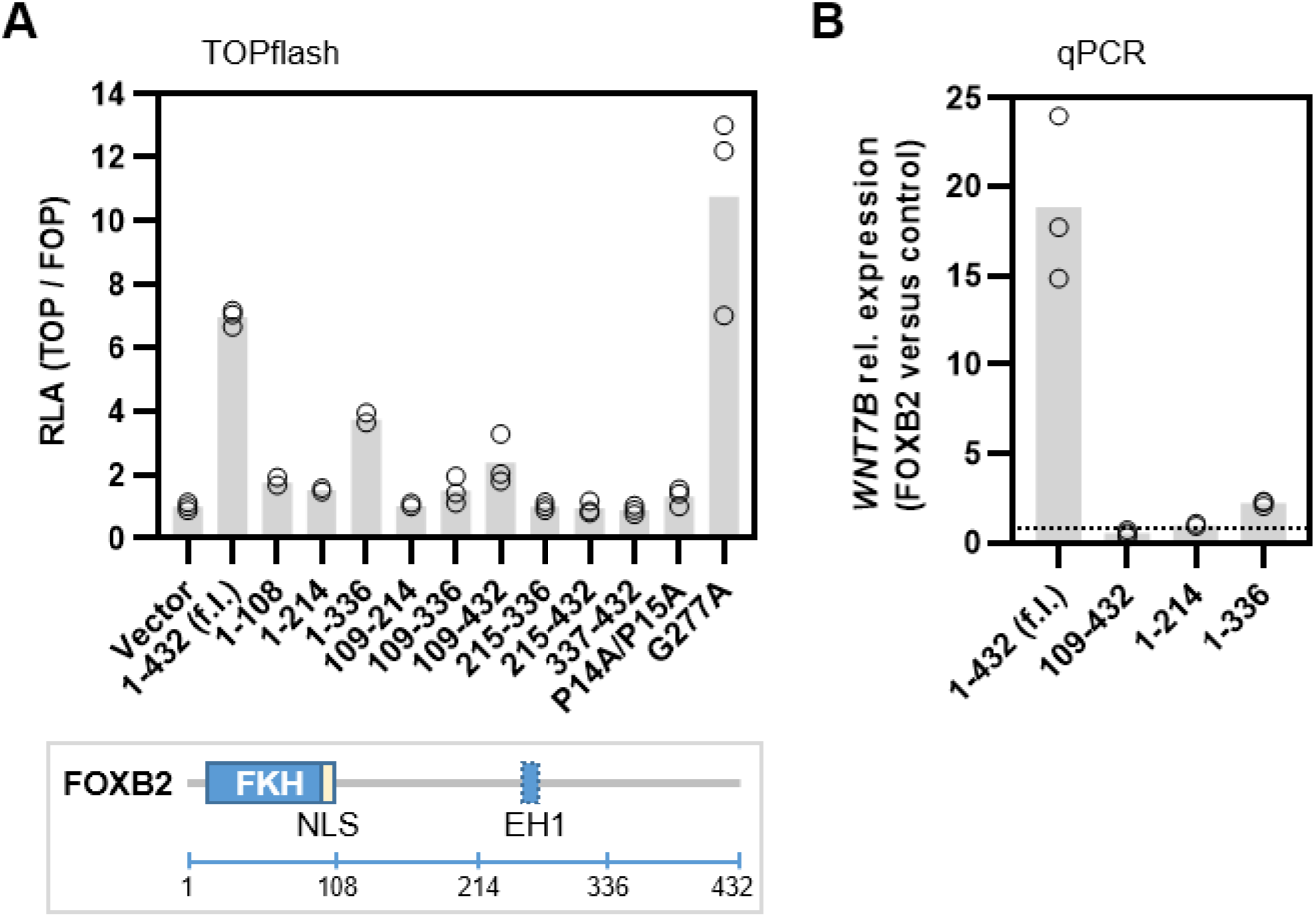
FOXB2-dependent Wnt signaling requires multiple protein domains. (**A**) FOXB2 mutation or truncation decreased its TOPflash activity in 293T cells. Numbers indicate amino acid positions. f.l., full length. Major protein features are shown in the cartoon below. FKH, forkhead domain; NLS, nuclear localization sequence; EH1, engrailed homology 1 motif. (**B**) qPCR analysis in 293T cells showed that FOXB2 N or C-terminal truncation blocked *WNT7B* induction.

### FOXB2-dependent Wnt signaling is controlled by transcriptional co-regulators

Our data so far suggested that activation of Wnt signaling by FOXB2 requires interaction with other proteins. In order to identify possible FOXB2 interactors, we generated an N-terminal BirA-FOXB2 fusion construct for proximity labeling (Roux et al., 2012), which retained full activity in TOPflash (Fig. S4B-D). Mass spectrometry following streptavidin pull-down of pulse-labeled proteins revealed numerous candidates that were enriched compared to free BirA control, with high consistency across multiple experiments (Fig. 4A). Statistical analysis narrowed the initial list down to 95 high confidence hits. These included several candidates that have previously been linked to Wnt signaling, such as YAP1 and JUN (Fig. 4B). As expected, gene ontology analysis of high confidence interactors showed that putative FOXB2-associated proteins are primarily involved in the regulation of gene transcription and mRNA splicing (Fig. 4C, and Supplemental Table 2). In contrast, and in agreement with our earlier observations, we did not pull down β-catenin in any experiment.

**Figure 4:**
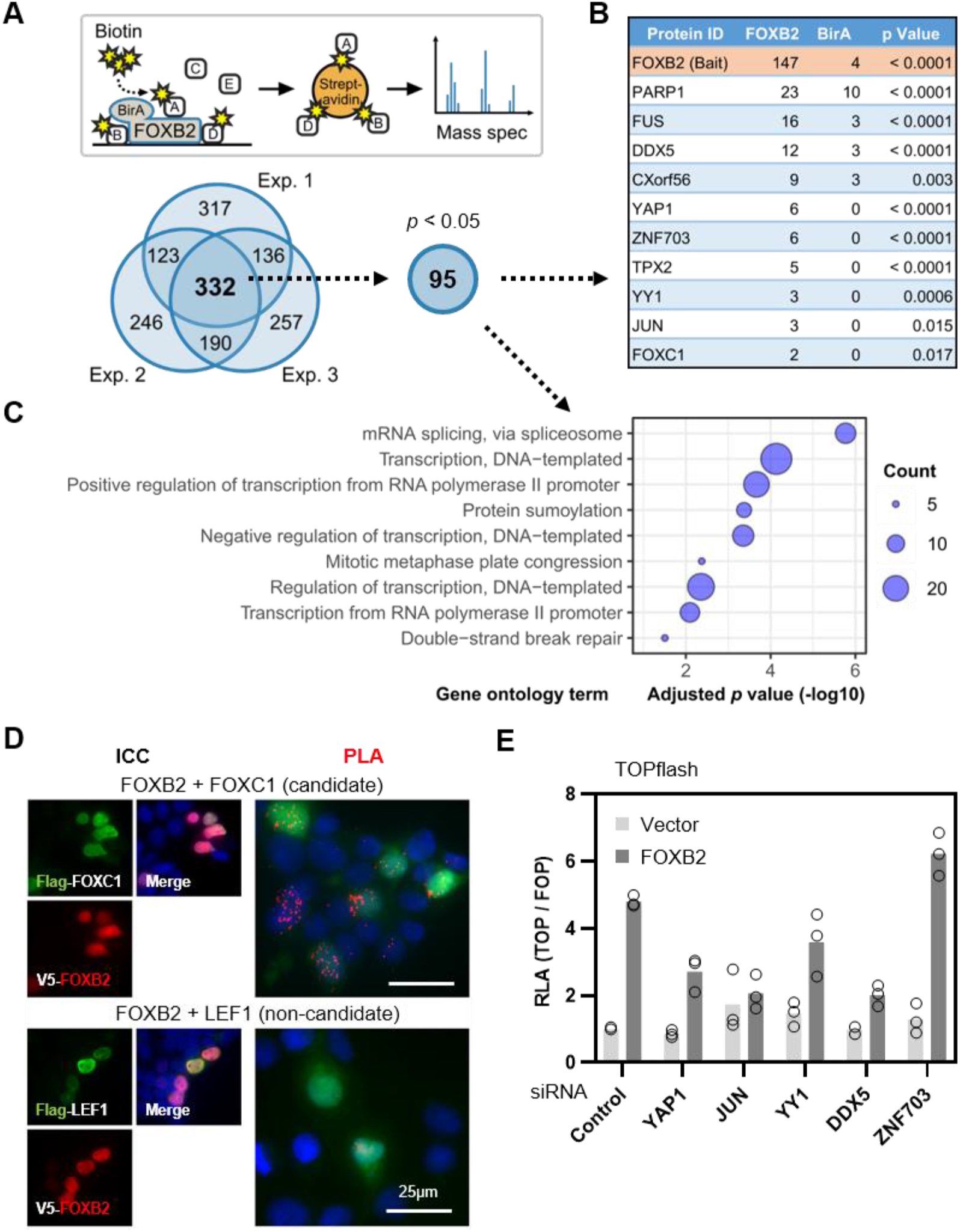
Transcription co-regulators contribute to FOXB2-dependent Wnt signaling. (**A**) Schematic summary of the BioID assay used to identify FOXB2 interactors. The Venn diagram below indicates the number of proteins identified in three independent BioID experiments performed in 293T cells, after subtraction of BirA background. High confidence FOXB2 interactors (Fisher’s exact test *p* < 0.05) were analyzed further. (**B**) Partial list of candidate interactors. Numbers indicate the average score from three experiments. (**C**) Gene ontology analysis of FOXB2 interactors. Only GO terms with an adjusted *p* < 0.05 are shown. (**D**) In-situ proximity ligation assay (PLA) in 293T cells, and corresponding immunocytochemistry (ICC). FOXC1, but not LEF1, interacted with FOXB2 in the nucleus, as indicated by red dots. (**E**) TOPflash assay in 293T following depletion of candidate FOXB2 interactors. Several co-regulators modulated FOXB2-dependent Wnt signaling. Note that these siRNAs had no effect on basal Wnt signaling.

We validated the BioID data by in-situ proximity ligation (Fig. 4D). Indeed, candidate interactor FOXC1, but not LEF1 control, was confirmed as a FOXB2-associated protein in situ. Moreover, we tested the functional role of candidate interactors in FOXB2-dependent Wnt signaling by RNA interference (Fig. 4E). We observed that several candidates, particularly YAP1, JUN, and DDX5, were required for FOXB2-dependent TOPflash activation, but they had no negative effect on basal Wnt signaling. In contrast, depletion of transcriptional co-repressor ZNF703 slightly increased FOXB2-induced Wnt pathway activation. We conclude that FOXB2 interacts with a multi-protein transcriptional complex to promote Wnt activity.

### FOXB2 is induced in aggressive prostate cancer

The only major expression site of mammalian FOXB2 is the developing brain, particularly the thalamus and hypothalamus (Kaestner et al., 1996). In adult mice, limited *Foxb2* expression was observed in some tissues, specifically the brain, thymus, ovary, and testis (Kaestner et al., 1993). Although analysis of public gene expression databases showed that *FOXB2* levels are exceedingly low in normal tissues, we found that it is induced in some cancers, including thymomas, ovarian cancer, and testicular germ cell cancer (Fig. S5). In particular, we observed that *FOXB2* transcript levels were frequently increased in prostate cancer. Here, FOXB2 amplification was detected mainly in aggressive, castration-resistant and neuroendocrine tumors (Fig. 5A). On the molecular level, highest *FOXB2* expression was observed in the iCluster 2 prostate cancer subtype (Cancer Genome Atlas Research, 2015), which is predominantly characterized by *ERG* fusions, as well as *PTEN* and *TP53* mutation (Fig. S6A). Of note, these genetic lesions are individually associated with prostate cancer progression and poor prognosis, further suggesting that FOXB2 is specifically induced in aggressive cancers.

**Figure 5:**
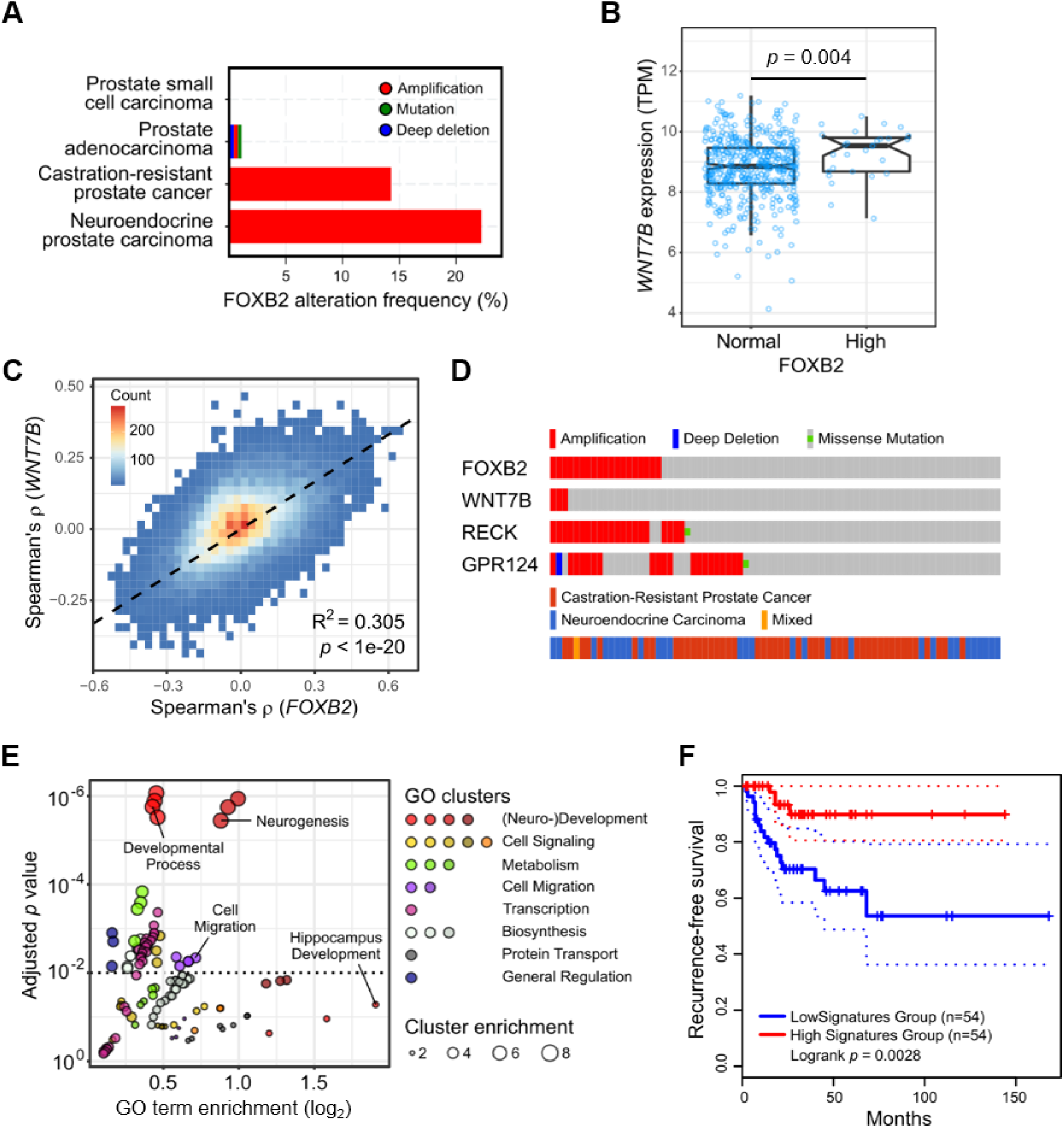
FOXB2 is induced in advanced prostate cancer. (**A**) FOXB2 genomic alterations across 14 prostate cancer studies curated in the cBioPortal for Cancer Genomics. FOXB2 is amplified specifically in advanced, castration-resistant and neuroendocrine cancers. (**B**) RNA-seq expression analysis in prostate adenocarcinoma (TCGA dataset PRAD) showed increased *WNT7B* levels in tumors with FOXB2 upregulation. (**C**) Genome-wide correlation density plot in dataset PRAD. Data analysis showed a high degree of correlation between the WNT7B and FOXB2-associated transcriptome. (**D**) OncoPrint analysis of aggressive prostate cancers (Beltran et al., 2016) revealed frequent co-amplification of FOXB2 and the WNT7B co-receptors RECK and GPR124, but not WNT7B itself. (**E**) Gene ontology (GO) analysis of the top 500 positively FOXB2/WNT7B-associated genes in prostate cancer. GO terms were clustered based on functional relation. Individual GO terms of interest are highlighted. (**F**) Kaplan-Meier recurrence-free survival plot in iCluster 2 prostate cancer, based on the top 50 FOXB2/WNT7B negatively correlated genes (see also Fig. S8). Dashed lines indicate the 95% confidence interval.

Subsequent data analysis revealed that *FOXB2* upregulation in prostate cancer was associated with significantly increased *WNT7B* levels (Fig. 5B), and that *WNT7B* transcript levels were highest in iCluster 2 tumors (Fig. S6B). Importantly, genome-wide analysis of prostate cancer RNA-seq data showed a high degree of correlation between the FOXB2 and WNT7B-associated transcriptome, suggesting that they act in the same pathway in cancer (Fig. 5C). Consistent with this notion, the WNT7B receptors RECK and GPR124, but not WNT7B itself, are frequently co-amplified with FOXB2 in castration-resistant and neuroendocrine prostate cancers (Fig. 5D). In contrast, FOXB1 and FOXA2 were not associated with increased *WNT7B* expression (Fig. S6C).

Gene ontology analysis of the most strongly FOXB2/WNT7B co-correlated genes in prostate cancer revealed that positively correlated genes are primarily involved in neurogenesis and cell migration (Fig. 5E), which is consistent with the anticipated physiological role of *Foxb2* in brain development (Kaestner et al., 1996). Conversely, negatively correlated genes mainly control small molecule metabolism and metal ion transport, required for normal prostate function (Fig. S6D). Taken together, these data suggest that FOXB2, presumably via WNT7B, may regulate a transcriptional program involved in the neuronal differentiation of prostate cancer cells, a feature that is linked to cancer progression and poor prognosis (Parimi et al., 2014).

In support of this hypothesis, we observed that both FOXB2 and WNT7B are associated with worse recurrence-free survival (RFS) in prostate cancer (Fig. S7A, B), albeit non-significantly in the case of FOXB2. Moreover, numerous FOXB2/WNT7B correlated genes are individually associated with altered RFS in prostate cancer, and remarkably, they cluster in a near-binary manner (Fig. S7C): genes that are positively correlated with FOXB2 preferentially predict worse survival, whereas negatively correlated genes are linked to better survival. Indeed, we observed that a gene signature consisting of the top 50 FOXB2/WNT7B negatively correlated genes was strongly associated with improved RFS in iCluster 2-type prostate cancer (Fig. 5F, and Fig. S7D, E), where *FOXB2* and *WNT7B* expression peaks. We conclude that the FOXB2/WNT7B-associated prostate cancer transcriptome promotes disease progression, by favoring malignant cell differentiation.

### FOXB2 drives LNCaP neuroendocrine differentiation

We tested this hypothesis in LNCaP prostate carcinoma cells, which undergo neuroendocrine differentiation in response to various stimuli, including Wnt pathway activation (Uysal-Onganer et al., 2010; Yang et al., 2005). We first confirmed regulation of Wnt signaling by FOXB2 in LNCaP cells (Fig. 6A). Although results were less consistent than in other cell lines, presumably due to low transfection efficiency, we observed robust activation of the TOPflash reporter by FOXB2, at least in the presence of exogenous Wnt3a and R-spondin 3.

**Figure 6:**
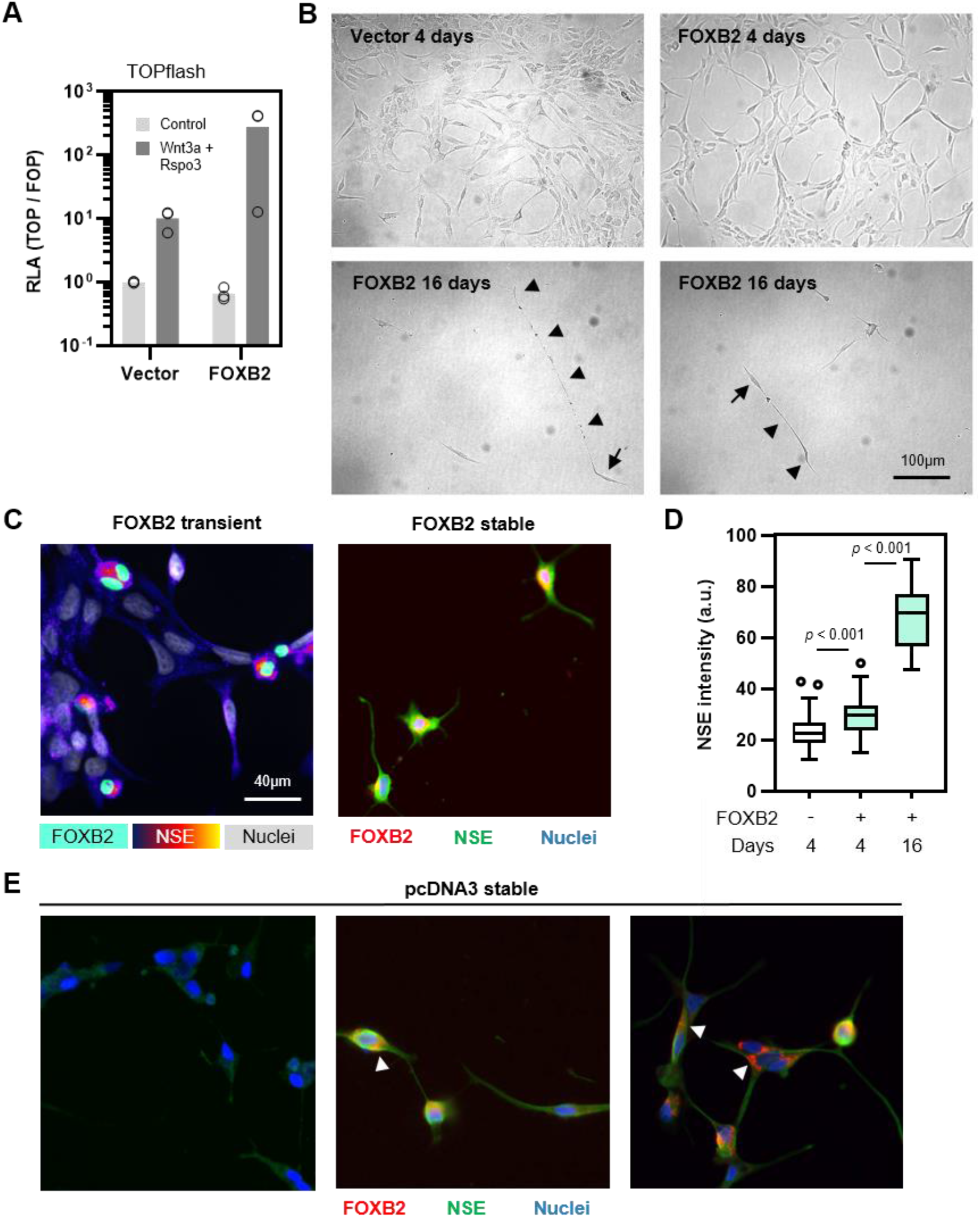
FOXB2 induces LNCaP neuroendocrine differentiation. (**A**) TOPflash assay in LNCaP prostate cancer cells. FOXB2 synergized with Wnt3a/Rspo3 in pathway activation. (**B**) Brightfield images of LNCaP cell transfected with FOXB2 or empty vector control for the indicated times. Arrows highlight individual cells with very long, dendrite-like processes (arrowheads). (**C**) Immunofluorescence staining of neuronal marker NSE in LNCaP cells. FOXB2 overexpressing cells exhibited higher levels of NSE intensity. (**D**) Quantification of NSE mean fluorescence intensity after transient (4 days) or stable (16 days) expression of FOXB2. Data were analyzed using Bonferroni’s multiple comparison test following analysis of variance (ANOVA). (**E**) FOXB2/NSE staining in pcDNA3 control transfected cells after G418 selection for > 2 weeks. Images show different staining patterns within the same sample. Arrowheads indicate extranuclear FOXB2 staining.

We next asked whether FOXB2 could cause prostate cancer neuroendocrine differentiation. Transient transfection of FOXB2 in LNCaP cells for up to 4 days did not induce any obvious phenotypical changes (Fig. 6B). However, stable FOXB2 expression resulted in a rapid, near-complete growth arrest following selection in G418-containing media, which precluded us from propagating these cells. Remaining cells adopted a neuron-like morphology with compact body and very long protrusions. Importantly, this phenotype is consistent with LNCaP neuroendocrine differentiation induced by e.g. androgen depletion and Wnt11 overexpression (Uysal-Onganer et al., 2010).

We validated this observation by investigating the expression of differentiation marker neuron-specific enolase (NSE) (Fig. 6C, D). After four days, FOXB2-expressing cells displayed slightly but significantly elevated NSE levels compared to non-transfected cells in the same sample. Moreover, stable expression of FOXB2 substantially increased NSE expression, consistent with neuroendocrine differentiation. Conversely, we found that in empty vector transfected control LNCaP subjected to the same G418 selection protocol, some cells underwent spontaneous differentiation, albeit at a lower rate than FOXB2 transfected cells (Fig. 6E). Further investigation revealed that transformed cells displayed elevated NSE levels, and some of them expressed endogenous FOXB2 protein, which was undetectable in any other assay or cell line we tested. Curiously, FOXB2 staining occasionally localized to perinuclear speckles or cell protrusions. Whether this pattern is caused by unspecific binding of our FOXB2 antibody, or indicates an extranuclear function of FOXB2, remains to be determined. Taken together, these data suggest that FOXB2 can be induced during prostate cancer progression, and is sufficient to drive the neuroendocrine differentiation of cancer cells.

## DISCUSSION

The key conclusions from our study are i) that FOXB2, an uncharacterized protein, is a potent regulator of Wnt ligand expression and TCF signaling, and ii) that FOXB2 drives neuroendocrine prostate cancer differentiation. Our results thus add FOXB2 to the growing list of FOX transcription factors involved in Wnt pathway regulation and pathobiology. FOX proteins comprise an evolutionarily conserved transcription factor family with 43 members in humans, which play critical roles in development and tissue homeostasis (Golson and Kaestner, 2016; Myatt and Lam, 2007). Consistently, dysregulation of FOX signaling is a common feature of major human diseases, notably cancer. Despite this, the function of many FOX proteins, including FOXB2, remains poorly understood.

FOX transcription factors affect tumorigenesis in part by controlling Wnt signaling, which is a key oncogenic pathway in various cancers (Nusse and Clevers, 2017). In particular, several FOX proteins directly engage the β-catenin signaling complex, and thereby regulate target gene expression. FOXM1 and FOXQ1, for example, control the nuclear translocation of β-catenin, and drive the progression of brain and colorectal cancer (Peng et al., 2015; Zhang et al., 2011). Similarly, FOXK1/2 activate Wnt signaling in colon cancer, by controlling the nuclear shuttling of scaffold protein Disheveled (Wang et al., 2015). Wnt signaling is also prominently involved in prostate cancer pathogenesis (Murillo-Garzón and Kypta, 2017), but it is unclear to what extent FOX transcription factors control Wnt activity in this disease.

Our results show that FOXB2 is induced during the neuroendocrine differentiation of LNCaP prostate carcinoma cells, and that its levels are increased in aggressive prostate cancer. Moreover, we find that FOXB2 activates the Wnt pathway via induction of agonistic Wnt ligands, primarily WNT7B. *WNT7B* expression in prostate cancer has been linked to tumor growth and drug resistance (Miyamoto et al., 2015; Zheng et al., 2013), and may thus contribute to malignant cancer progression. Interestingly, Zheng et al. reported that *WNT7B* is a direct transcriptional target of the androgen receptor (AR) (Zheng et al., 2013). AR expression is frequently lost in advanced prostate cancer, particularly neuroendocrine prostate cancer, but *WNT7B* expression is actually highest in androgen-independent prostate carcinoma cells (Bonkhoff, 2001; Zhu et al., 2004). FOXB2 induction may thus maintain high WNT7B levels in the absence of androgen signaling, and thereby drive cancer cell differentiation and treatment resistance. Moreover, Zheng et al. suggested that WNT7B promotes prostate cancer via protein kinase C signaling, rather than TCF/LEF (Zheng et al., 2013). Whether FOXB2 also acts through non-canonical Wnt pathway(s), especially in prostate cancer, remains to be determined. However, both canonical and non-canonical Wnt signaling modes have been implicated in prostate cancer neuroendocrine differentiation (Uysal-Onganer et al., 2010; Yang et al., 2005), and FOXB2 may engage both pathways in parallel through induction of multiple Wnt ligands. Given the unresolved signaling mode(s) of WNT7B, especially also in Wnt/TCF signaling (Alok et al., 2017), unraveling its role in prostate cancer may have broader implications for Wnt signaling in general.

Besides its role in Wnt ligand regulation, we find that FOXB2 engages numerous β-catenin nuclear interactors, including FUS, NONO, PARP1, DDX5, TOP2A, and hnRNP K (Sato et al., 2005), but not β-catenin itself. Moreover, we also consistently observed association of TCF4 with FOXB2 in the BioID assay, although these results did not meet our criteria for statistical significance. Given the observation that FOXB2 can induce *AXIN2* in β-catenin-deficient cells, we consider it possible that FOXB2 may coopt the β-catenin transcriptional machinery, to independently drive the expression of TCF target genes. In this context, it is interesting to note that Iee et al. recently identified TCF4 as a major determinant of prostate cancer neuroendocrine differentiation (lee et al., 2019). Thus, FOXB2 may additionally act as a “rogue” transcription (co)-factor that activates AR and TCF signaling in a ligand-independent manner; however, this hypothesis remains to be tested more rigorously.

With regard to developmental biology, prior reports suggest that there is substantial overlap between *Foxb2, Wnt7b*, and *Axin2* expression domains in the developing brain, particularly in the early thalamus (Bluske et al., 2009; Kaestner et al., 1996; Shimogori et al., 2010). During embryogenesis, WNT7B signaling is primarily involved in angiogenesis and the formation of the blood-brain barrier, which it controls redundantly with WNT7A (Stenman et al., 2008). Consequently, *Wnt7a/Wnt7b* double knock-out mice die around E12.5 due to fatal cerebral hemorrhaging. We find that FOXB2 strongly induces *WNT7B*, and to a lesser extent *WNT7A*, at least in 293T cells. Thus, FOXB2 may be a rheostat of Wnt signaling activity and angiogenesis in the developing brain.

In summary, we report the first molecular characterization of the cryptic forkhead box transcription factor FOXB2. We identify FOXB2 as a potent activator of Wnt/TCF signaling in normal and cancer cells. Given the putative roles of FOXB2 in neurogenesis and the neuroendocrine differentiation of advanced prostate cancer, it is possible that FOXB2 controls both processes through a common, presumably WNT7B-dependent signaling mode, which is aberrantly re-activated in cancer. We believe that further exploration of the FOXB2/WNT7B-associated transcriptome may uncover new therapeutic vulnerabilities in rare, aggressive cancers.

## ACKNOWLEDGEMENTS

The authors thank Drs Christof Niehrs, Claudio Cantù, and Lennart Svensson for cell lines and reagents. We also thank all investigators who have made materials and data available through public repositories. Technical support from the microscopy and mass spectrometry core facilities at Linköping University is gratefully acknowledged. S.K. is a Wallenberg Molecular Medicine fellow, and receives financial support from the Knut and Alice Wallenberg Foundation.

## AUTHOR CONTRIBUTIONS

L.M. designed, performed, and analyzed most experiments with help from G.P. S.K. conceived and supervised the study, performed experiments, analyzed experimental and public data, and wrote the manuscript with input from all authors.

## SUPPLEMENTAL FIGURES

**Figure S1:**
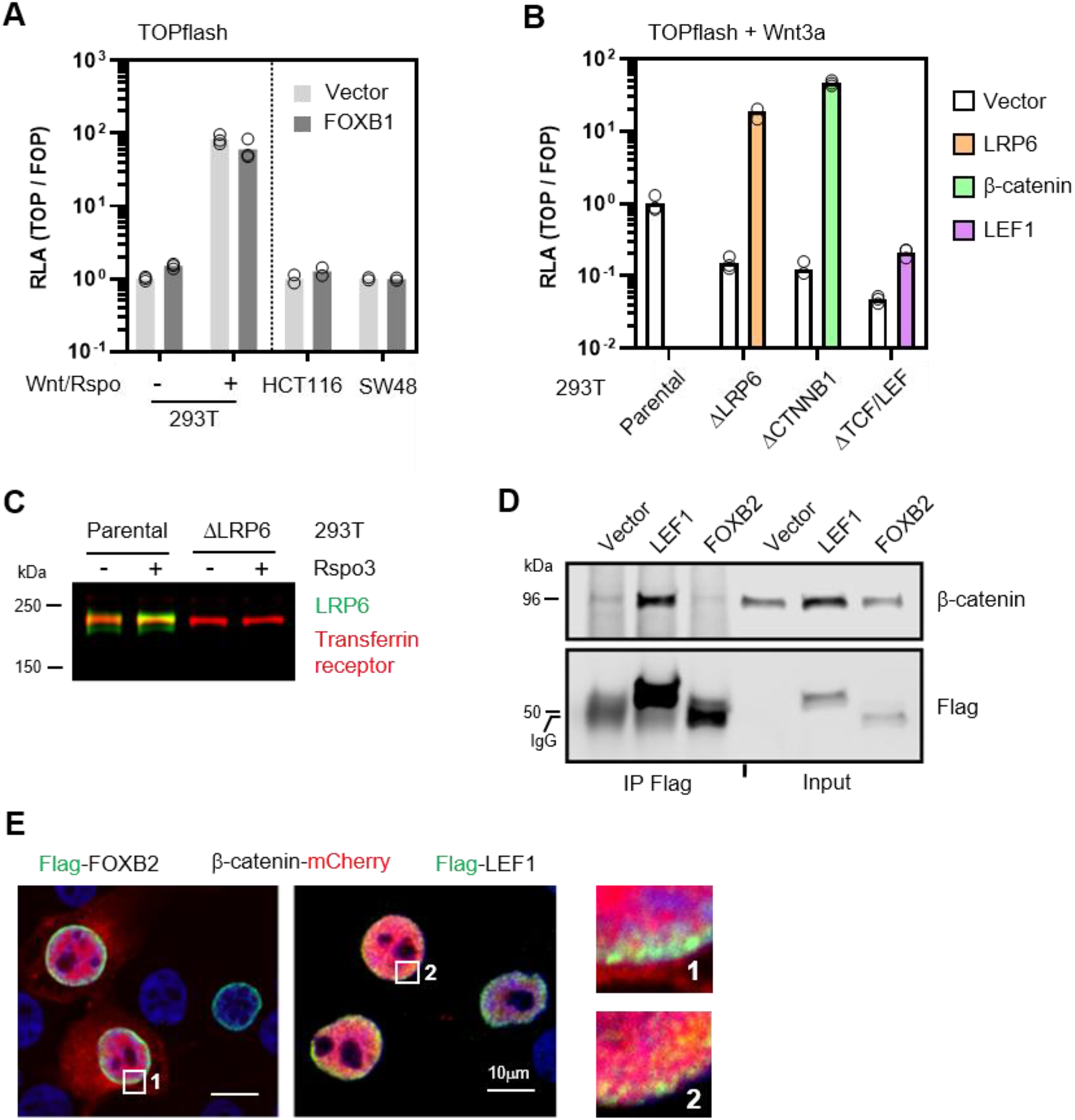
FOXB2 activates Wnt signaling upstream of β-catenin. (**A**) TOPflash assay in the indicated cell lines showed that FOXB1 does not regulate Wnt/β-catenin signaling. (**B**) TOPflash assay in the presence of exogenous Wnt3a showed strongly attenuated reporter activity in gene-edited 293T cells. Re-expression of the missing pathway component rescued their loss-of-function. (**C**) Immunoblot for LRP6 confirmed loss of Wnt receptor expression on the protein level in ΔLRP6 cells used in reporter assays. (**D**) Co-immunoprecipitation from nuclear lysates of HCT116 cells. The indicated Flag-tagged proteins were pulled down using anti-Flag antibodies, and endogenous β-catenin was detected by immunoblotting. LEF1, but not FOXB2, associated with β-catenin. (**E**) Co-expression of FOXB2 and LEF1 with β-catenin in 293T cells. β-catenin co-localized with LEF1, but not FOXB2, as indicated by the yellow signal. Note also the lack of β-catenin nuclear focusing in FOXB2-expressing cells.

**Figure S2:**
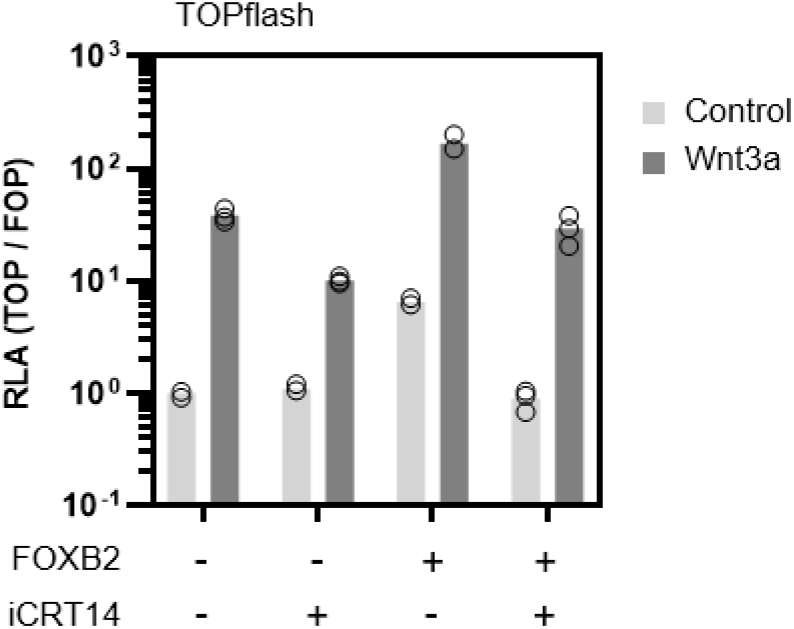
FOXB2-dependent Wnt activity requires β-catenin/TCF interaction. TOPflash assay in the presence of 10μM iCRT14 showed attenuated pathway activation by FOXB2.

**Figure S3:**
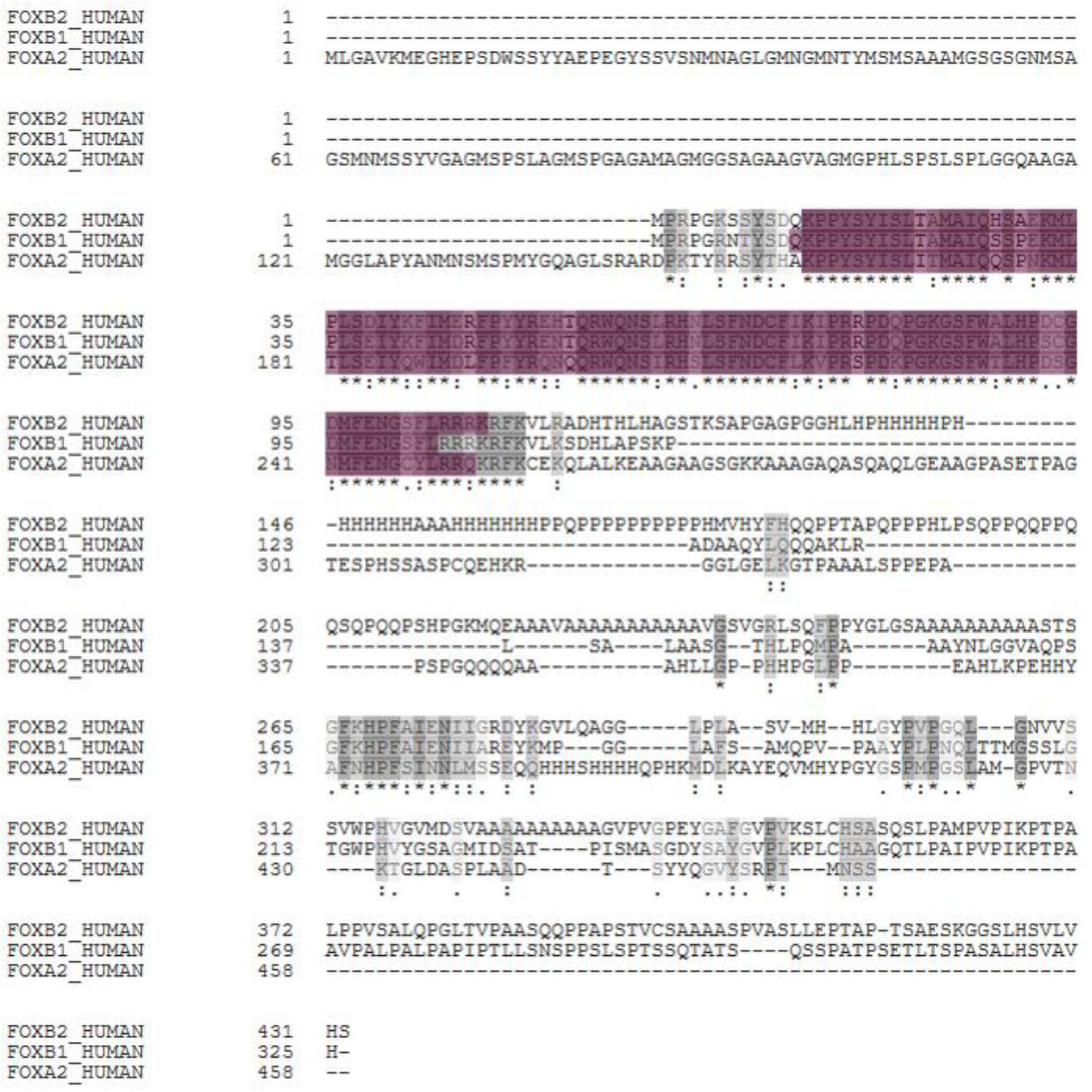
The forkhead box is highly conserved in FOX proteins. Sequence alignment of human FOXB2 (Accession Q5VYV0), FOXB1 (Q99853), and FOXA2 (Q9Y261). The DNA-binding forkhead box is highlighted in pink. Sequence similarity is indicated by shades of grey.

**Figure S4:**
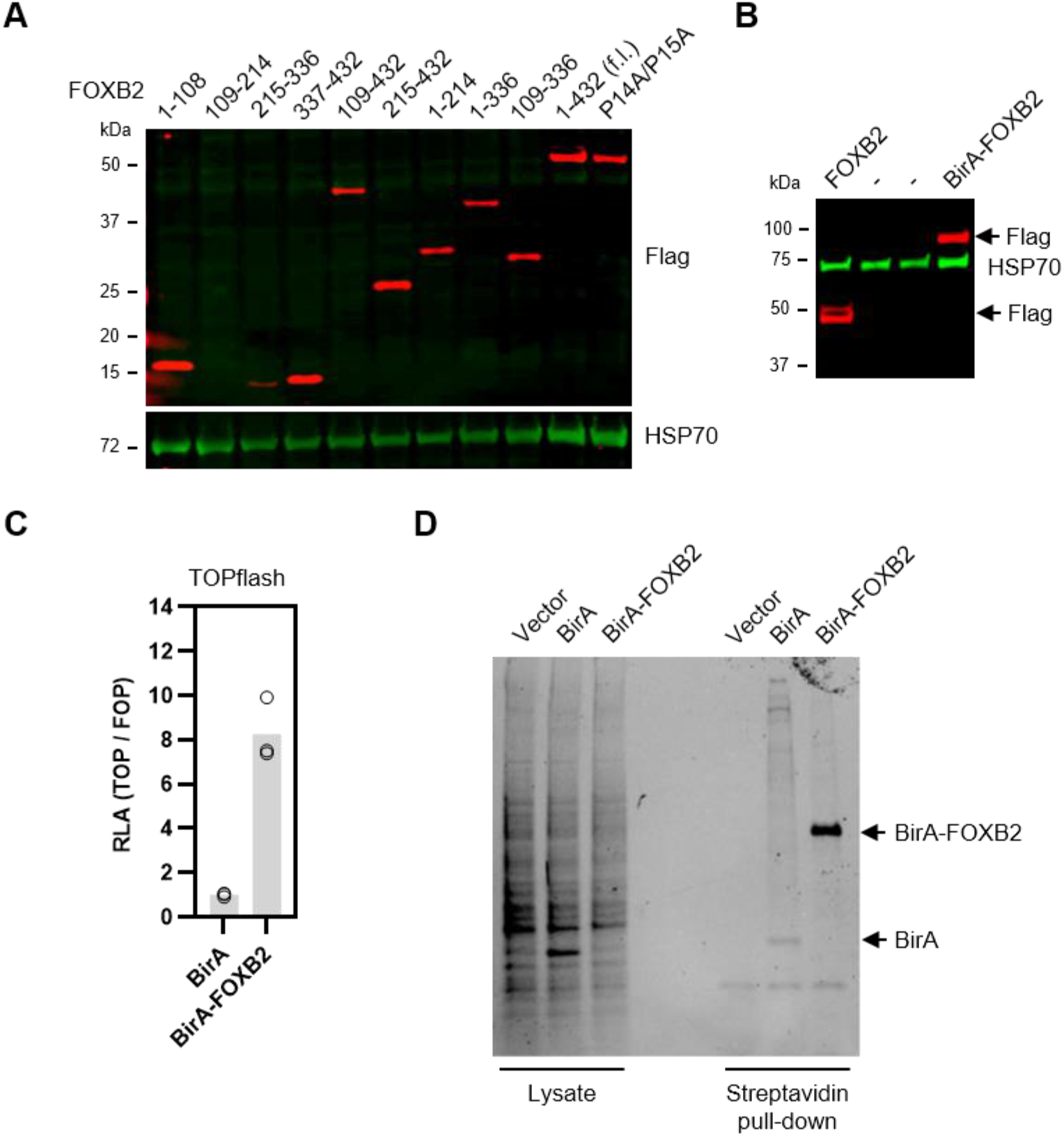
Analysis of FOXB2 constructs. (**A, B**) Protein expression analysis of Flag-tagged FOXB2 mutants. Note that construct 109-214 was poorly expressed in this, but not subsequent experiments. (**C**) TOPflash assay in 293T cells. Note that data are from the same assay as in Fig. 3A, and were normalized to empty vector control. (**D**) Streptavidin pull-down of biotinylated proteins in 293T cells, following BirA or BirA-FOXB2 expression and pulse-labeling. Proteins were visualized by UV activation of TGX stain-free gels (Bio-Rad).

**Figure S5:**
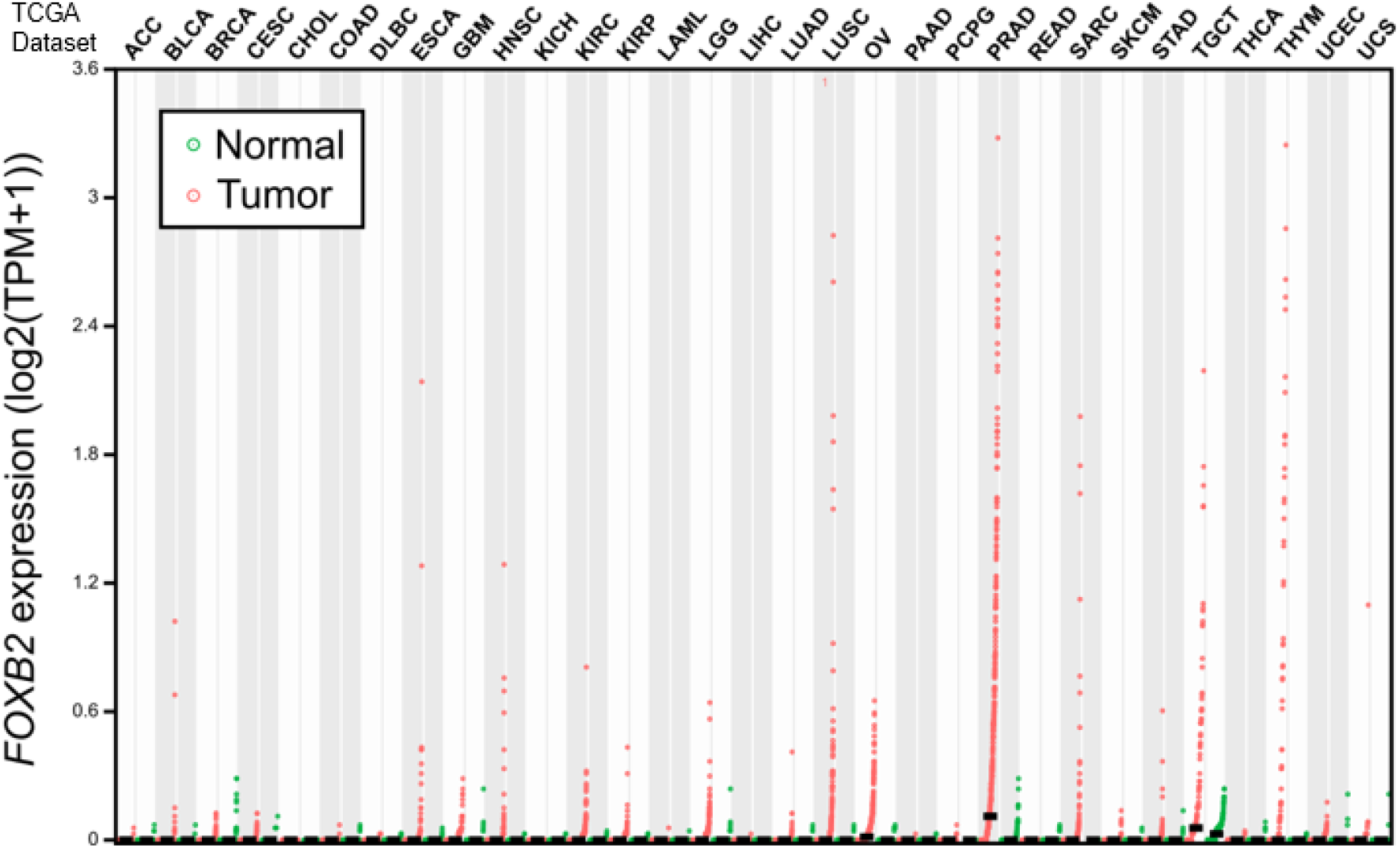
FOXB2 is induced in select cancers. RNA-seq analysis showed *FOXB2* induction in select cancer types. Cancer data from the Cancer Genome Atlas (TCGA) were matched to available TCGA normal expression data from the same tissue. PRAD, prostate adenocarcinoma; OV, ovarian serous cystadenocarcinoma; THYM, thymoma. Explanation of other cancer types can be found on the TCGA homepage at https://cancergenome.nih.gov/.

**Figure S6:**
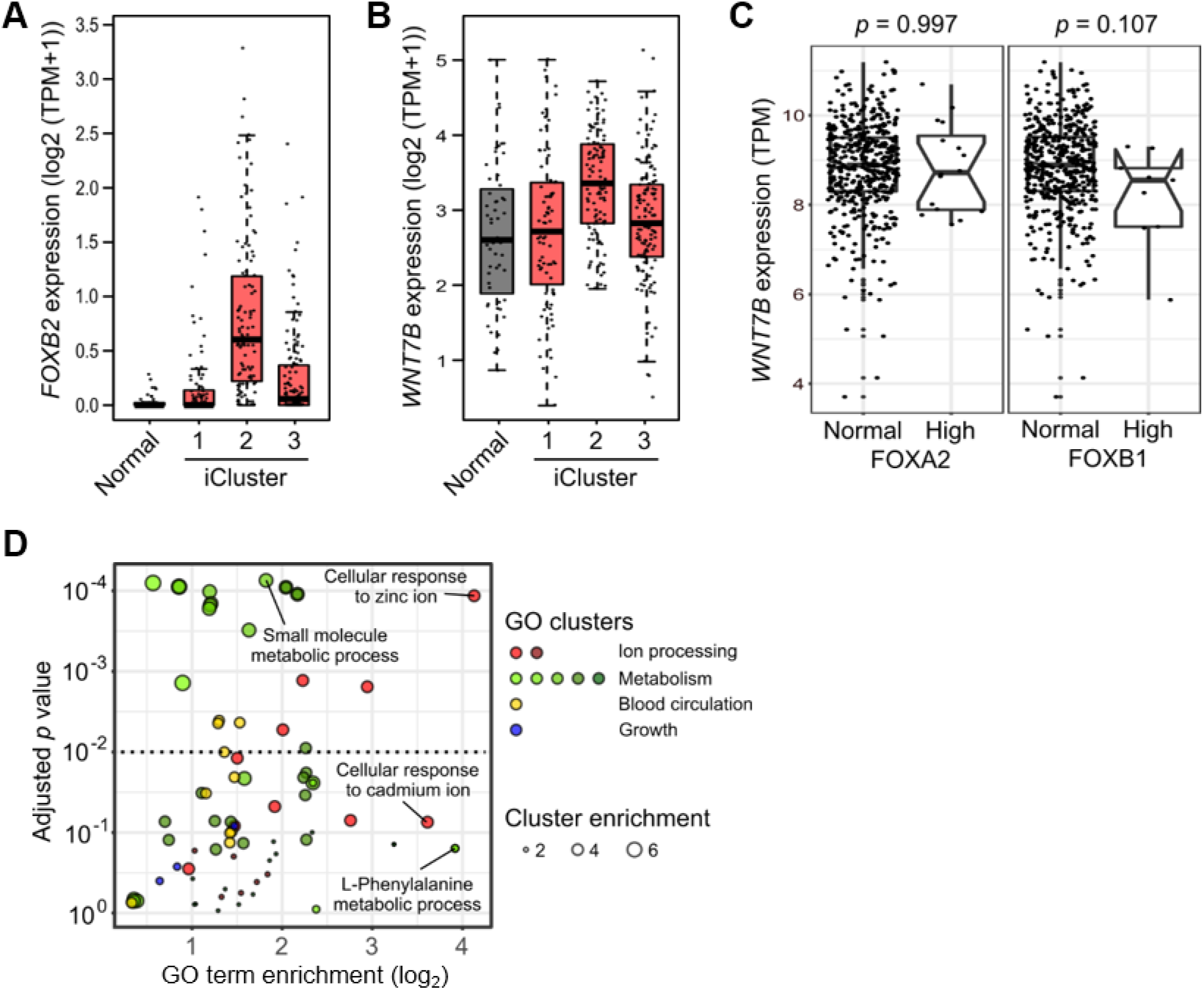
FOXB2 is associated with WNT7B in advanced prostate cancer. RNA-seq expression analysis of (**A**) *FOXB2* and (**B**) *WNT7B* in normal prostate and prostate cancer. Tumors were stratified by iCluster molecular subtype, as described in (Cancer Genome Atlas Research, 2015). *FOXB2* and *WNT7B* levels were concomitantly elevated in the aggressive iCluster 2 subtype. (**C**) RNA-seq analysis of TCGA dataset PRAD showed no association between *FOXA2/WNT7B* and *FOXB1/WNT7B* levels. (**D**) Gene ontology (GO) analysis of the top 500 negatively FOXB2/WNT7B-associated genes in prostate cancer. GO terms were clustered based on functional relation. Individual GO terms of interest are highlighted.

**Figure S7:**
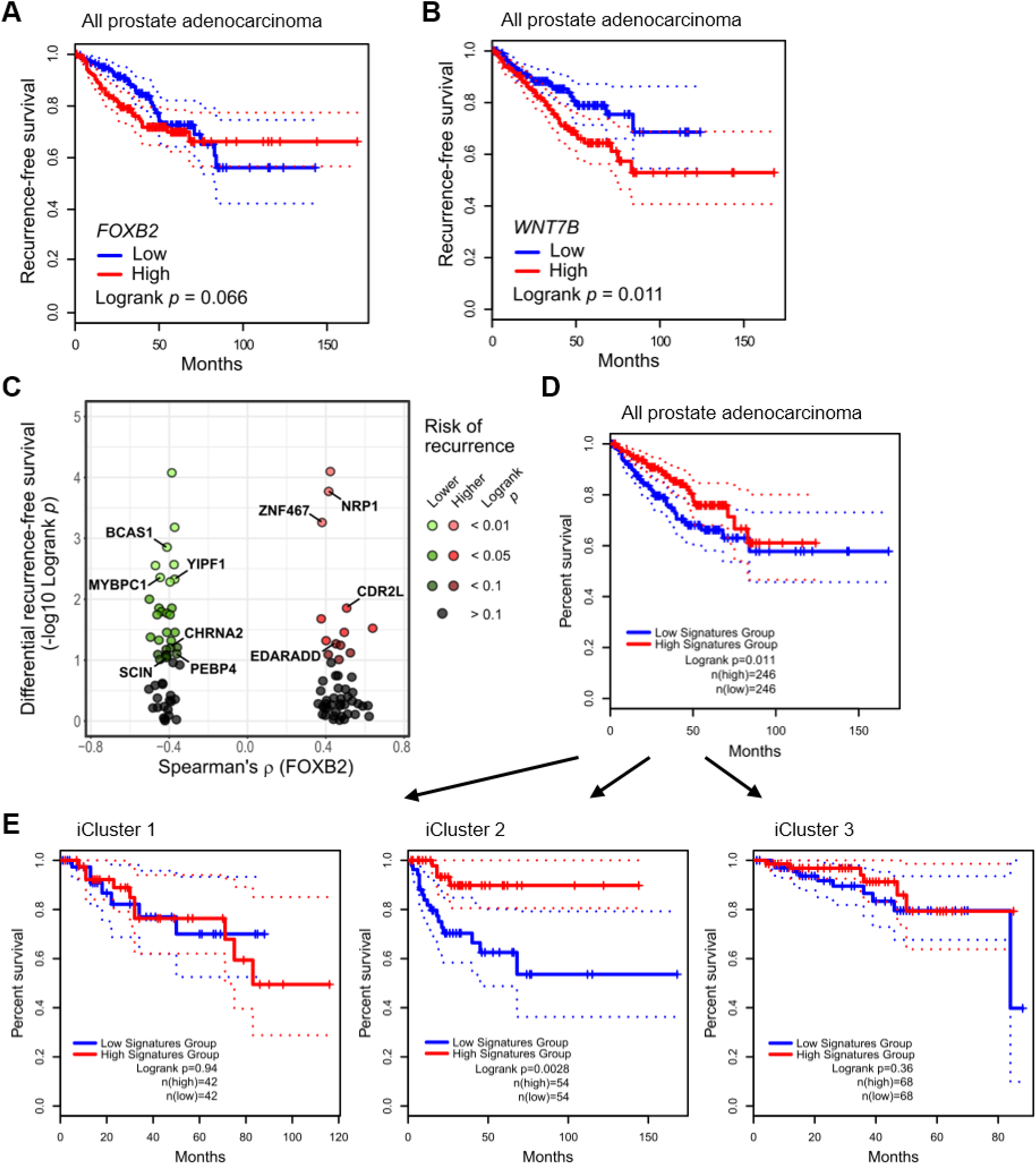
FOXB2 and WNT7B are associated with poor prognosis in prostate cancer. Kaplan-Meier plot of recurrence-free survival in TCGA dataset PRAD, based on expression of (**A**) *FOXB2* and (**B**) *WNT7B*. Both genes were associated with unfavorable prognosis. (**C**) In-depth survival analysis of the top 50 positively and negatively FOXB2/WNT7B-associated genes in prostate cancer. The plot shows the Spearman correlation value of each gene against FOXB2 in dataset PRAD, versus their individual prognostic value in recurrence-free prostate cancer survival. Some genes of interest are highlighted. (**D, E**) Further analysis of the top 50 negatively correlated genes (left half in panel C) in recurrence-free prostate cancer survival. The 50 gene signature was associated with better prognosis in all prostate cancer (**D**), and particularly in the iCluster 2 subtype (**E**).

## SUPPLEMENTAL TABLES

**Supplemental Table 1:** Plasmids used in this study.

| Construct | Backbone | Addgene # | Reference |
| --- | --- | --- | --- |
| Flag-FOXB2 | pCS2+ N-Flag | - | This study |
| V5-FOXB2 | pCS2+ N-V5 | - | This study |
| Flag-FOXB1 | pCS2+ N-Flag | - | This study |
| Flag-FOXQ1 | pCS2+ N-Flag | - | This study |
| Flag-FOXF2 | pCS2+ N-Flag | - | This study |
| Flag-FOXC1 | pCS2+ N-Flag | - | This study |
| Flag-LEF1 | pCS2+ N-Flag | - | This study |
| FOXB2 | pcDNA3 | - | This study |
| Flag-FOXB2 1-108 | pCS2+ N-Flag | - | This study |
| Flag-FOXB2 1-214 | pCS2+ N-Flag | - | This study |
| Flag-FOXB2 1-336 | pCS2+ N-Flag | - | This study |
| Flag-FOXB2 109-214 | pCS2+ N-Flag | - | This study |
| Flag-FOXB2 109-336 | pCS2+ N-Flag | - | This study |
| Flag-FOXB2 109-432 | pCS2+ N-Flag | - | This study |
| Flag-FOXB2 215-336 | pCS2+ N-Flag | - | This study |
| Flag-FOXB2 215-432 | pCS2+ N-Flag | - | This study |
| Flag-FOXB2 337-432 | pCS2+ N-Flag | - | This study |
| Flag-FOXB2 P14A/P15A | pCS2+ N-Flag | - | This study |
| Flag-FOXB2 G277A | pCS2+ N-Flag | - | This study |
| WNT7B reporter | pTA-Luc | - | This study |
| BirA | pCS2+ N-Flag | - | This study |
| BirA-Flag-FOXB2 | pCS2+ N-Flag | - | This study |
| M50 Super 8x TOPFlash | pTA-Luc | 12456 | (Veeman et al., 2003) |
| M51 Super 8x FOPFlash | pTA-Luc | 12456 | (Veeman et al., 2003) |
| HA-LRP6 | pCS2+ N-HA | - | Christof Niehrs, German Cancer Research Center |
| Flag-Rspo3ΔC | pCS2+ N-Flag | - | (Ohkawara et al., 2011) |
| mCherry-Beta-Catenin-20 | mCherry | 55001 | Michael Davidson, Florida State University |
| eSpCas9(1.1) | pX330-U6-Chimeric_BB-CBh-hSpCas9 | 71814 | (Slaymaker et al., 2016) |
| eGFP | pcDNA3 | - | Peter Zygmunt, Lund University |

**Supplemental Table 2**, containing high confidence FOXB2 interactors and their associated gene ontology analysis, can be found in the online supplemental material.

